# Behavioral and Neuroanatomical Investigation of Attention Deficit Hyperactivity Disorder Pathogenesis in Juvenile Spontaneously Hypertensive Rats

**DOI:** 10.1101/2023.01.04.522787

**Authors:** Aysegul Gungor Aydin, Esat Adiguzel

## Abstract

Attention-deficit/hyperactivity disorder (ADHD) is one of the most prevalent neuropsychiatric disorders of childhood, characterized by locomotor hyperactivity, impaired sustained attention, impulsivity, and distractibility. Recently, the dysfunction of different synaptic circuits in the prefrontal cortex (PFC) has been shown. Previous studies have attributed the pathophysiological mechanism of ADHD to disturbances in the dopaminergic system. In this study, we tested the hypothesis that spontaneously hypertensive rats (SHR), which are considered a validated animal model of ADHD, have altered dopaminergic innervation and increased locomotor activity. Here, we performed immunohistochemical tyrosine hydroxylase (TH) and dopamine-beta-hydroxylase (DBH) staining. The mesocortical dopaminergic system appears to be normal in juvenile SHR, as suggested by (i) no alteration in the area density of TH-immunoreactive (TH-ir) dopaminergic neurons in the VTA, (ii) no alterations in the volume density of TH-ir fibers in layer I of the PrL subregion of mPFC, (iii) no alteration in the percentage of TH-ir dopaminergic fibers in layer I of the PrL subregion of mPFC as revealed by TH and/or DBH immunoreactivity. Furthermore, the SHR showed increased locomotor activity than WKY in the open field test.

The demonstration of no alteration in mesocortical dopaminergic neurons and fiber in SHR raises some concern about the position of SHR as an animal model of the inattentive subtype of ADHD. However, these results strengthen this strain as an animal model of hyperactive/impulsive subtype ADHD for future studies that may elucidate the underlying mechanism mediating hyperactivity and test various treatment strategies.

## 1. INTRODUCTION

Attention-deficit hyperactivity disorder (ADHD) is one of the most prevalent neuropsychiatric disorders of childhood, characterized by locomotor hyperactivity, impaired sustained attention, impulsivity, and distractibility (American Psychiatric Association, 2013). The worldwide prevalence of ADHD is 5.29%, also the prevalence of childhood and adulthood ADHD is 5-10% and 4%, respectively (Faraone *et al*., 2003; Kessler *et al*., 2006; Polanczyk *et al*., 2007). Multiple hypotheses have been proposed for the etiology of ADHD (Biederman *et al*., 1995; Giedd *et al*., 2001; Kreppner *et al*., 2001; Mick *et al*., 2002; Nigg & Casey, 2005), but one that has stood the test of time is the dopamine (DA) deficit theory (Swanson et al., 2007). Previous studies have attributed the pathophysiological mechanism of ADHD to disturbances in the dopaminergic system (Volkow et al., 2007; Bowton et al., 2010).

There is a significant amount of data in the literature suggesting a DA-ADHD association. It is reported that the alterations in DA signaling in both individuals (Jucaite *et al*., 2005; Ludolph *et al*., 2008; Volkow *et al*., 2009) and rodents (Carr & White, 1987; Canales & Iversen, 1998; Somkuwar et al., 2016) can be correlated to ADHD symptoms. ADHD may result from deficits in the dopaminergic system, especially the mesocortical one, which projects from the VTA to the PFC (Sullivan & Brake, 2003; Robbins & Arnsten, 2009). The mesocortical DA inputs exert an inhibitory effect on the activity of PFC neurons (Sesack & Bunney, 1989; Pirot *et al*., 1992). Due to its strong inhibitory effect on prefrontal cortical neurons, dysfunction of the mesocortical dopaminergic projections is involved in the behavioral problems of children with ADHD (Pirot *et al*., 1992). Animal studies showed that lesions of the mesocortical projection to PFC lead to behavioral problems similar to those seen in ADHD (Bubser & Schmidt, 1990).

The medial prefrontal cortex (mPFC) of the rat is most likely comparable to that of humans (Papa *et al*., 2002; Heidbreder & Groenewegen, 2003; Kelly *et al*., 2007). Neurochemical, pharmacological, genetic, and imaging studies in human and animal models highlight the critical role of catecholamine dysfunction and particularly DA in cortical brain structures such as the PFC in the neurobiology of ADHD (Prince, 2008; Smith *et al*., 2008; Fernández *et al*., 2009; Naneix *et al*., 2009; Arnsten, 2011; Cortese *et al*., 2012; Moon *et al*., 2014; Qin *et al*., 2016). The PFC requires optimal levels of catecholamine neurotransmitters norepinephrine (NE) and DA for regulating PFC-dependent executive functions that are often reported to be suboptimal in ADHD patients (Xing *et al*., 2016). Hence, attentional, psychomotor, reinforcing, and rewarding behaviors in which DA plays an essential modulatory role are deficient in ADHD (Brozoski *et al*., 1979; Himelstein *et al*., 2000).

The Spontaneously hypertensive rat (SHR) is currently the best-validated animal model of ADHD based on behavioral, genetic, and neurobiological data (Sagvolden & Johansen, 2012). SHR shows overactivity under the control of a fixed-interval operant reinforcement schedule (Berger & Sagvolden, 1998; Sagvolden, 2000), increased behavioral variability (Mook *et al*., 1993), and problems with cognitive impulsiveness (Evenden, 1998) similar to that of ADHD children. Abnormalities in DAT-1 gene expression are seen both in ADHD individuals (Cook *et al*., 1995; Mill *et al*., 2005) and in the SHR (Russell *et al*., 1998). Furthermore, chronic oral methylphenidate at therapeutically relevant doses improves behavioral and cognitive deficits in SHR (Kantak *et al*., 2008; Harvey *et al*., 2013). In addition, SHR exhibits hyperactivity, as seen in ADHD children, in the open field test (Tsai et al., 2017).

Several studies have suggested that either too little, or too much D1 receptor stimulation impairs PFC function (Sawaguchi & Goldman-Rakic, 1994; Murphy *et al*., 1996; Zahrt *et al*., 1997; Granon *et al*., 2000). Clinical and experimental evidence suggests the involvement of DA systems, especially the mesocortical systems, in ADHD (Sullivan & Brake, 2003). The mesocortical DA pathway, originating from the ventral tegmental area (VTA) and projecting to the PFC, is involved in cognitive functioning (Floresco & Magyar, 2006; Castner & Williams, 2007). However, it is still unclear whether the mesocortical dopaminergic system is hyper or hypo-functioning. Therefore, in the present study, we tested the hypothesis that SHR show hypofunctional dopaminergic neurotransmission. We addressed this hypothesis by setting out to determine whether the DA neurons and/or axons deficiency might be the cause of the postulated dopaminergic hypofunction in SHR. Thus, immunohistochemical analysis was carried out on neurons and fibers of mesocortical DA systems of the SHR and WKY. WKY rats are considered a proper control for the SHR as they were both established from some paternal, normotensive Wistar stock (Sagvolden & Johansen, 2012). Also, our study aimed to analyze the dopaminergic neurons and fibers in SHR and WKY to search for neuroanatomical evidence that might explain the hypofunctional dopaminergic system and assess the face validity of this animal model in the open field test.

## 2. MATERIALS AND METHODS

### 2.1 Animals

Juvenile (4-week-old) male Spontaneously Hypertensive Rats (SHR) (n=15) and Wistar Kyoto rats (WKY) (n=13) (50–60 g on arrival, Charles River Laboratories, Wilmington MA) were used in the present study. Out of the total, 20 (SHR: n=10; WKY: n=10) and 8 rats (SHR: n=5; WKY: n=3) were used in behavioral and anatomical procedures, respectively. All animals were housed in a colony room with a 12-hour light/dark cycle (light on at 8.00 am) and had ad libitum access to water and standard rat feed. All anatomical procedures were approved by the animal care and use committee of the University of Virginia and under the National Institutes of Health guidelines. The behavioral experiments were performed at Pamukkale University and under the approval of 2015/09. As common in SHR literature (Botanas et al., 2016; Somkuwar et al., 2016; Cheng et al., 2017), only male rats were used in this study since ADHD-like symptoms are more prevalent in male rats (Biederman, 2005), and ADHD is three times more prevalent in males than females (Barkley, 1990; Swanson *et al*., 2001). Moreover, the same rat strain had behavioral testing in the present study which proved one of the main ADHD symptoms (increased locomotor activity) in SHR in the open field task. Furthermore, we measured the systolic blood pressure of animals prior to experiments to rule out the possible confounding factor i.e. hypertension.

### 2.2 Blood pressure measurements

Animals had been trained to stay in the rat holder to condition the animal for this procedure which was repeated in three days and were tested at the beginning of the experiment. Noninvasive systolic blood pressure (SBP) measures obtained from day 4 were considered valid. Conscious rats were restrained in a warming chamber at 34°C (RXRESTRAINER-S; BIOPAC BioPac Systems) for 10–20 min before non-invasive SBP measurements using a computerized indirect tail-cuff method (NIBP200A, BioPac Systems). The sensor (RXTCUFSENSOR9.5-9.5mm; BioPac Systems), consisting of an infrared light source and infrared light detectors mounted in a 95 mm long inflatable rubber cuff, was placed on the base of the tail of the rats then SBP was recorded (Lehnen *et al*., 2010). The mean SBP is an average of four measurements made over a 10-min period.

### 2.3 Behavioral testing

All the animals, SHR (n=10) and WKY(n=10), were removed from the colony room and brought to the behavioral testing area in their home cages 30 mins before the behavioral testing to minimize the stress due to exposure to a new environment. Behavioral testing was performed in a quiet room at the beginning of the light phase of the light/dark cycle. At the end of the test, the number of fecal boluses was counted, and the arena was cleaned with 70% ethanol prior to use to remove any scent clues left by the previous animal.

#### 2.3.1 Locomotor activities in open field test

To validate the face validity of SHR, the animal was placed in the center, always facing the same direction, and allowed to explore the open field for 10 min. The locomotor activity was assessed using a computerized and automated activity monitoring apparatus (May Act 508, 42×42× 42 cm; Commat Ltd, Ankara, Turkey) capable of tracking different behavioral activities. The apparatus comprised a black floor divided into 196 equal squares and surrounded by a transparent wall equipped with infrared sensors. Multiple variables were measured (distance moved, immobility, horizontal, vertical, and stereotypy movements).

### 2.4 Tissue Preparation and Sampling Design

The processing of tissue from each of these eight rats used for anatomical studies was as follows: sections from 8 brains (5 SHR, 3 WKY) were used for single immunolabeling for TH, light microscopy, and sections from 6 brains (3 SHR and 3 WKY) were used for dual immunolabeling for dopamine DBH and TH, confocal microscopy. The TH-ir was considered to be the most optimal marker to identify dopaminergic neurons (Bayer & Pickel, 1990) and fibers (Hokfelt *et al*., 1976; Sesack *et al*., 1998). However, because cortical noradrenergic fibers also express TH (Pickel *et al*., 1976), DBH immunoreactivity is widely utilized as a specific biomarker for cortical noradrenergic fibers (Morrison *et al*., 1978). We employed double-labeling techniques with anti-TH and anti-DBH antibodies to determine whether the TH-ir fibers do indeed selectively label dopaminergic fibers in mPFC.

The rats were deeply anesthetized with an overdose of euthosole (excess of 150 mg/kg, i.p) and transcardially perfused using room-temperature Tyrode’s solution (137 mM NaCl, 5.5 mM Dextrose/Glucose, 1.2 mM MgCl_2_, 2 mM KCl, 0.4 mM NaH_2_PO_4_, 0.9 mM CaCl_2_, 11.9 mM NaHCO_3_, in 1 L filtered dH_2_0) followed by 4% paraformaldehyde in 0.1 M phosphate buffer (PB; pH 7.4). Brains were removed and allowed to postfix overnight in 4% paraformaldehyde at 4°C. Four sets of coronal vibratome sections through the mPFC and VTA cut at 50 μm were collected in PB. All morphological analyses were done in sections obtained through the rostrocaudal extent, between Bregma 4.20 and 2.76 for mPFC and Bregma -5.52 and -6.00 for VTA, according to Stereotaxic Rat Atlas (Paxinos, G.& Watson, 2007). Adjacent sections were mounted on glass slides and stained with Nissl to determine the borders of the region of interest. Sections that were not processed immediately were treated with 1% sodium borohydride (NaBH_4_; 1 g NaBH_4_ in 100 ml 0.1 M PB) to stop fixation, rinsed until the bubbles had cleared and stored in 0.05% sodium azide in phosphate-buffered saline (PBS) at 4°C.

### 2.5 Immunocytochemistry

Tissue sections were collected in PBS and blocked in 1% bovine serum albumin (BSA; Sigma Aldrich) in 0.01 M PBS, pH 7.4, for 30 minutes at room temperature. Free-floating tissue sections were then transferred in a chicken polyclonal antibody against TH (Abcam, Cat# AB76442, 1:1000; Table 1) and 1% BSA in 0.01 M PBS with 0.05% NaN_3_ and 0.3% Triton X-100 (Sigma Aldrich) for incubation of 1–3 days. The sections were then rinsed in 0.01 M PBS and transferred into biotinylated goat anti-chicken secondary antibody (Vector Laboratories, Burlingame, CA; Cat# BA-9010; 1:100 dilution) for 2 hours, followed by ABC-DAB visualization. All sections were mounted serially on gelatin-subbed slides, dehydrated, and coverslipped using DPX mounting media (Sigma Aldrich, St. Louis, MO).

**Table 1.**
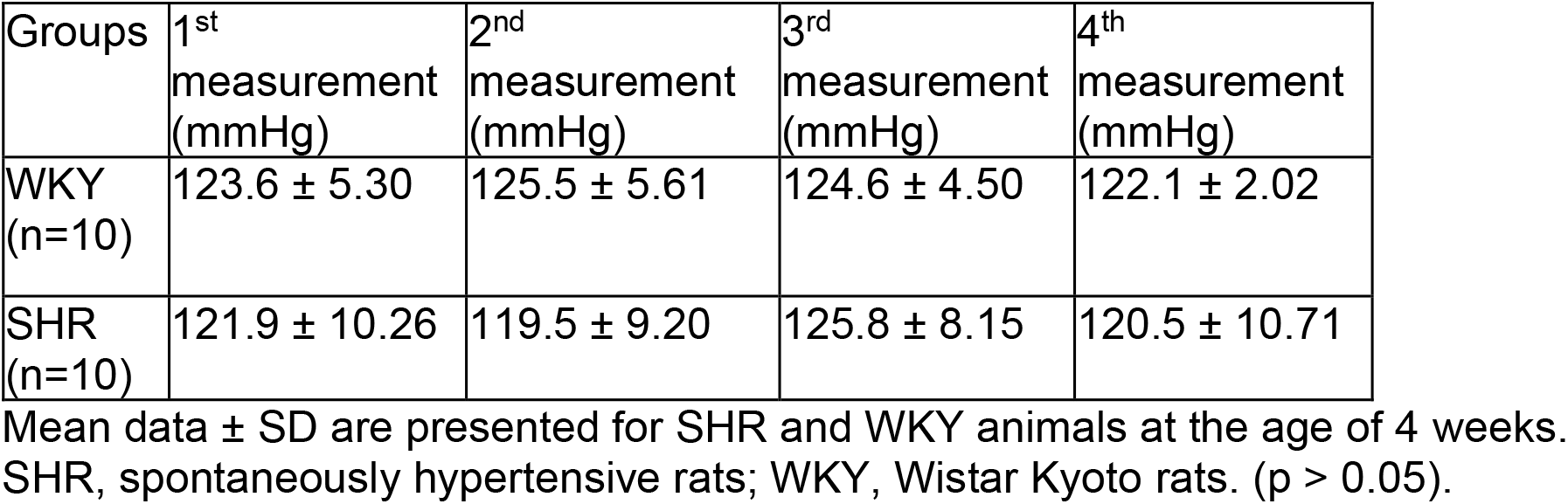
Systolic blood pressure from WKY and SHR rats. The table shows four different SBP measurements from the groups.

| Groups | 1<sup>st</sup><br>measurement<br>(mmHg) | 2<sup>nd</sup><br>measurement<br>(mmHg) | 3<sup>rd</sup><br>measurement<br>(mmHg) | 4<sup>th</sup><br>measurement<br>(mmHg) |
| --- | --- | --- | --- | --- |
| WKY<br>(n=10) | 123.6 ± 5.30 | 125.5 ± 5.61 | 124.6 ± 4.50 | 122.1 ± 2.02 |
| SHR<br>(n=10) | 121.9 ± 10.26 | 119.5 ± 9.20 | 125.8 ± 8.15 | 120.5 ± 10.71 |

Mean data ± SD are presented for SHR and WKY animals at the age of 4 weeks. SHR, spontaneously hypertensive rats; WKY, Wistar Kyoto rats. (p > 0.05).

**Table 2.** List of antibodies and dilutions used in the study. The table summarizes all antibodies used and their species specificity, dilutions, and sources.

| Name | Host | Antigen characteristics | Source | Dilution |
| --- | --- | --- | --- | --- |
| Anti-Tyrosine hydroxylase | Chicken (polyclonal) |  | Abcam, AB76442; | 1:1000 |
| Anti-Dopamine Beta hydroxylase | Mouse (monoclonal) | Clone 4F10.2 (Milstein 2007) | Millipore Co. Temecula, CA, MAB308 | 1:300 |
| Anti-Chicken secondary antibody | Goat | Chicken IgG, conjugated to biotin | Vector Laboratories, Burlingame, CA; BA-9010 | 1:100 |
| Anti-Mouse secondary antibody | Donkey | Mouse IgG, conjugated to rhodamine (TRITC) | Jackson ImmunoResearch Laboratories, Inc., West Grove, PA; #715-025-151 | 1:500 |
| Anti-Chicken secondary antibody | Donkey (polyclonal) | Chicken IgG, conjugated to Alexa Fluor 488 | Jackson ImmunoResearch Laboratories, Inc., West Grove, PA, #: 703-545-155 | 1:500 |

Sections for dual-labeling with two markers were incubated in a blocking solution containing 1% bovine serum albumin in PBS for 30 minutes. The sections were then transferred into an antibody cocktail that contained 1:300 monoclonal anti-DBH (Millipore Co. Temecula, CA, catalog no MAB308; 1:300) and 1:1000 chicken polyclonal anti-TH (Abcam, Cat# AB76442, 1:1000), 0.05% NaN_3,_ 0.3 % Triton X-100, 1% BSA in PBS, and they were incubated at room temperature for 48 hours. Then the sections were rinsed three times in PBS (5 min each), and incubated in a mixture of donkey anti-mouse secondary antibody conjugated with rhodamine (TRITC) (red, Jackson Immunoresearch Laboratories, Inc., West Grove, PA; catalog no: 715-025-151,1:500) and polyclonal donkey anti-chicken secondary antibody conjugated with Alexa Fluor 488 (green, Jackson Immunoresearch Laboratories, Inc., West Grove, PA, catalog no: 703-545-155, 1:500) in PBS for 2 hours at room temperature in the dark. They were then rinsed three times in PBS and mounted on gelatin-coated glass slides, air-dried, and covered with an anti-fading mounting medium (Vector Laboratories, Burlingame, CA; catalog no. H-1000). The brain sections from two experimental groups were processed at the same time using the same preparations of the incubation solutions to eliminate variability due to protocol handling.

### 2.6 Image Acquisition

The sections processed for light microscopy were examined on a Leica DMLB microscope equipped with a digital camera (Leica MC170 HD). Low magnification images were taken with the 4x, and 10x objective lens, and high magnification images were taken with a 40x objective lens for quantitative analysis of TH-ir structures. The images were then adjusted for contrast and exposure in Adobe Photoshop. Architectural borders of VTA and layer I of the prelimbic (PrL) subregion of the mPFC (ROI) were identified after matching tissue sections with comparable atlas sections. These borders were also superimposed on adjacent sections that were stained for TH. Histoarchitectural transitions in the VTA and PrL subregion of the mPFC were examined in TH-stained sections. We obtained counts from both hemispheres, and all data for every analysis were pooled.

#### 2.6.1 Area Density of TH-ir neurons in VTA

Quantitative data were collected by one experimenter blinded to the animal experimental status using an unbiased sampling approach: We measured the volume density of TH-ir dopaminergic neurons in the VTA, using the Stereo Investigator (MBF Bioscience, Inc.) software. For this, the coronal sections that contained VTA were identified for analysis; these sections were comparable to plates # 79-82 of the rat brain atlas (Paxinos, G.& Watson, 2007). One-fourth of the sections from each brain were selected in accordance with systematic random sampling protocols, and four sections were used for counting cells (West *et al*., 1991). Using the ‘outline’ tool of the Stereo Investigator, the contour of the VTA was drawn under low magnification (4x objective). The selected areas were similar for all animals. Then, the outlined region was overlaid with a random series of counting frames (Sterio, 1984; Gundersen & Jensen, 1987). The TH-ir cell nuclei counts were performed under high magnification (40X). The top and bottom 5 µm at the top and bottom of the 50 μm thick sections were considered exclusion zones to reduce prevent over or underestimation related to surface artifacts. When the nucleus of a TH-ir neuron was unambiguous within the counting frame (50×50μm), it was included in the cell count. In a range of 53 and 89 counting frames in four sections were evaluated per each animal. Each counting frame consisted of two exclusion lines (left and bottom edges) and two inclusion lines (right and top edges). TH-ir cell nuclei were counted if they were found entirely within a counting frame or transected by at least one of the inclusion lines of a counting frame but not any of the exclusion lines of the same counting frame (Gundersen, 1977). The area TH-ir dopaminergic neuron density (N_A_) in the ROI was calculated using the following formula: N_V_ = N / A_ROI_ where N is the total number of TH-ir neurons and A_ROI_ is the total area analyzed (number of counting frame x area of counting frame) (Gundersen & Jensen, 1987; Witelson *et al*., 1995).

#### 2.6.2 Fiber Density Analysis using Brightfield Microscope

In order to quantify the density of TH-ir fibers, we outlined the PrL subregion of the mPFC using landmarks derived from Paxinos and Watson (Paxinos, G.& Watson, 2007). The ROI was outlined under low magnification (4×) and a 150μm wide measurement frame is positioned perpendicular to the cortical surface, at 10x magnification. Six different measurement frames per animal were used. Using a 40X objective, each TH-ir fiber in the ROI was traced, and the total length of fibers was computed using Neurolucida® software (MBF Bioscience, Inc.). The area of the measurement frame was also computed. The fiber density (μm/μm_3_) in each measurement frame was determined by dividing the total TH-ir fiber length over the measurement frame box volume. The volume of the measurement frame box on each image was obtained by multiplying the area of the measurement frame box with the thickness of the section.

#### 2.6.3 Fiber Density Analysis using Confocal Microscopy

Fibers were imaged using an 80i microscope fitted with a C2 scanning system (Nikon Instruments, Inc., Melville, NY) and a 10× objective (Nikon, CFIPlanApo; NA = 0.45). The fibers were matched for the wavelengths of the two lasers in the system (argon laser, 488 nm, 10 mW, Alexa Flour 488; DPSS laser, 561 nm, 10 mW, TRITC). A 60x objective (Nikon, PlanApo VC; NA =1.4) was used with a multi-track scanning method to completely separate the detection of the Alexa 488 and TRITC signals. A total of 3 SHR and 3 WKY animals were used and 9 images per animal were quantified. More precisely, 3 consecutive sections were imaged for the PrL subregion of the mPFC, and 3 images were acquired per section. All images were taken in lamina I. To avoid experimental bias, the experimenter was blinded to the experimental groups. For each image acquisition, the pixel size (0.21 μm), dwell time (10.8 μm), step size (0.25 μm), and z slice (5 μm) were the same. Each confocal image corresponded to a field of 1024×1024 pixels Photomicrographs for figures were made from maximum intensity images of the confocal stacks of single physical sections and were exported as a TIFF file to be analyzed with an image analysis system (Image-Pro Plus 7, Media Cybernetics, Inc, Silver Spring, MD). Only adjustments of brightness and contrast were used for images used in photomicrographs. The total length of TH-ir and DBH-ir fibers were evaluated in ROI with Image-Pro Plus 7 (Media Cybernetics, Inc, Silver Spring, MD) using manual tracing. The percentage of TH-ir fibers that were not also DBH-ir was determined by dividing the total TH-ir fiber length over the total fiber length.

### 2.7 Statistical Analysis

Data analysis was performed using Microsoft Excel 2007and and GraphPad Prism (GraphPad Software, San Diego, CA, USA) software. All graphs and figures were created using Prism (9.3.1; GraphPad Software, San Diego, CA, USA) and Adobe Photoshop CS5 (23.2.1), respectively. Descriptive data were obtained by using descriptive statistical methods such as mean, standard deviation, and median. SBP measurement and behavior data were submitted to the Shapiro–Wilk normality test, and the data displayed a normal distribution. For analysis of SBP data and open field data (distance moved, immobility, horizontal, vertical, and stereotypy movements) was conducted using an unpaired t-test. For immunohistochemical analysis, to compare the density of TH-ir neurons and fibers between groups, we used the Nested-t test. All statistical tests are two-tailed at a significance level of 0.05. Data on graphs represent mean ± standard deviation (SD). For all figures, ***p< 0.001, **p< 0.015, *p <0.05.

## 3. RESULTS

### 3.1 Systolic blood pressure

To test whether or not SHR is hypertensive SBP measurements are presented in Table 1. Non-invasive SBP measurement confirmed normotension at 4 weeks of age in the SHR group (121.9 ± 7.43 mmHg) and in the age- and sex-matched WKY group (124 ± 2.57 mmHg). The results revealed that there was no significant difference in SBP between the SHR (n=10) and WKY (n=10) groups (P > 0.05) (Fig 1).

**Fig 1.**
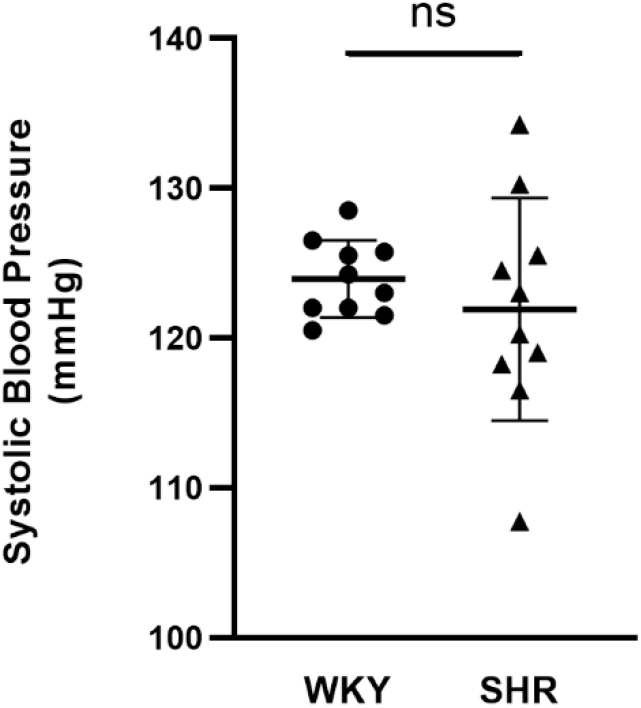
Tail cuff systolic blood pressure measurements of the WKY and SHR (n=10) animals at the age of 4 weeks. Values are expressed as mean ± SD. (p> 0.05, two-tailed unpaired t-test). No difference in SBP of SHR groups from corresponding WKY groups. SBP, systolic blood pressure.

### 3.2 Open field test

To assess the locomotor activity in SHR, an open field test was performed. Ten rats per strain were tested in this experiment. When placed into the open-field arena for the first time, i.e. non-habituated, SHR rats were significantly more active compared to WKY rats. The results are shown in Fig. 2, SHR rats showed significantly more horizontal (Fig. 2A, p<0.0001, t=5.574) and vertical activity (Fig. 2B p<0.0001, t=7.495), increased distance traveled (Fig. 2C p<0.0001, t=6.118), higher ambulatory activity (Fig. 2D p<0.001, t=4.689), and less immobility time (Fig. 2E p<0.01, t=3.650) than WKY rats. This finding indicates that SHR rats exhibited hyperactivity as revealed by the Open field test.

**Fig 2.**
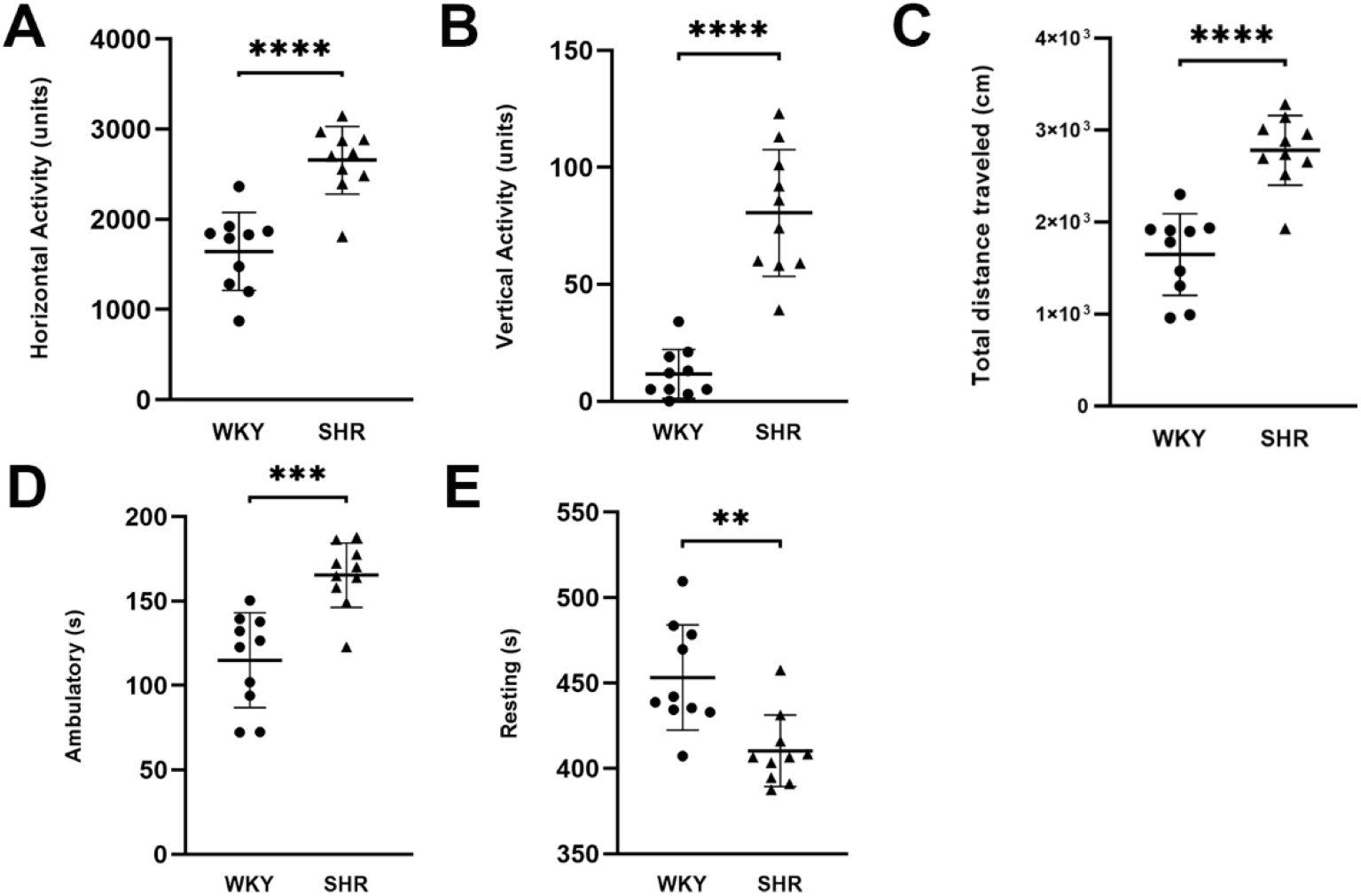
Increased locomotor activity in SHR rats compared to WKY rats in the open field test (n = 10 in each group). **(A)** Horizontal activity (arbitrary units), **(B)** Vertical activity (arbitrary units), (**C)** Total distance traveled (cm), (D) Ambulatory (s), (E) Resting (s). All data are presented as mean ± SD. SHR and WKY rats were compared with an independent sample T-test. A p-value <0.05 was considered significant. **p <0.01, ***p <0.001, ****p < 0.0001.

### 3.3 Area Density of TH-ir Dopaminergic Neurons in VTA

To assess whether there are fewer TH-ir neurons in VTA in the SHR, we measured the area density of TH-ir dopaminergic neurons. Fig. 1). The mean area density of neurons in the VTA was 6.2 ±2.9(x10^−3^) cell per mm_2_ (mean ±SD) for the control group, 5.7±2.8 (x10^−3^) cell per mm_2_ for the SHR group. The difference between the control and SHR groups was not statistically significant (p>0.05 from the Nested t-test).

### 3.4 Volume Density of TH-ir Fibers in layer I of the PrL subregion of mPFC

To determine whether the SHR has decreased TH-ir fibers in mPFC, we calculated the volume density of TH-ir fibers. The mean volume density of TH-ir fibers in layer I of the PrL subregion of mPFC was 0.0036 ± 0.0009 µm/µm_3_ (mean ±SD) for the control group, 0.0029 ± 0.0010 µm/µm_3_ for the SHR group (Fig. 2). The mean volume density of TH-ir fibers was significantly lower in SHR (p>0.05 from Nested t-test).

### 3.5 Percentage of TH-ir Dopaminergic Fibers in the mPFC

Because the TH-ir represents not only dopaminergic but also noradrenergic fibers from locus cereolous, we used dual; staining with TH and DBH antibodies to reveal the proportion of TH fibers that are not noradrenergic. Double immunofluorescent staining and colocalization analysis was performed on three slices from 3 areas of 3 animals. The percent ratio of TH-ir fibers that are not DBH+ (thus, dopaminergic) in layer I of the PrL subregion of mPFC (Fig. 3). This ratio in PrL subregion of mPFC in WKY (n=3) and SHR animals (n=3) were nearly equivalent 30.86 ± 6.07 (%) (mean ±SD) and 30.13 ±9.23 (%), respectively. There was no significant difference in the percentage of TH-ir dopaminergic fibers between WKY and SHR (p>0.05 from the Nested t-test).

**Figure 3.**
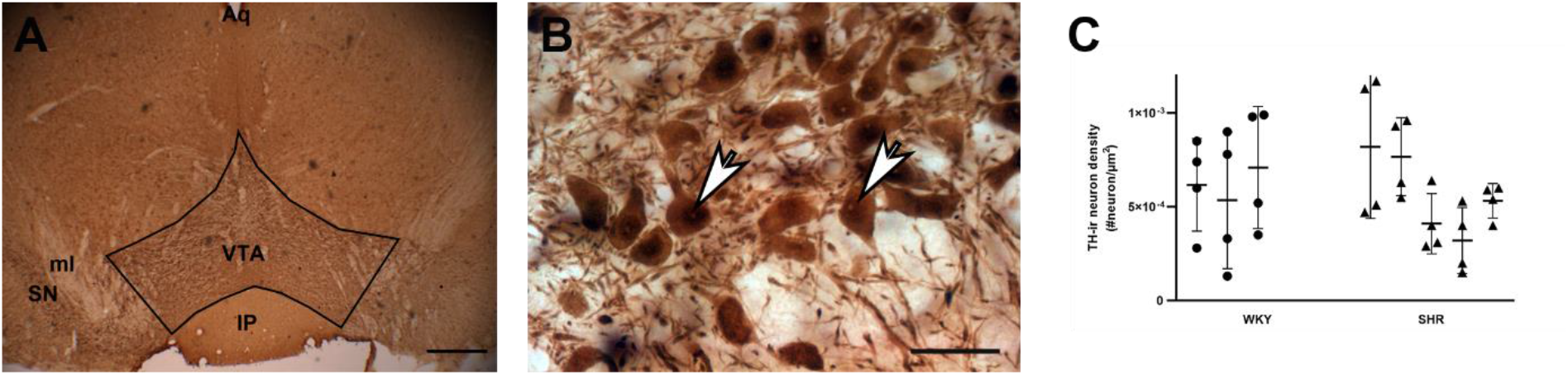
TH-ir neuron density in the VTA. (A) A coronal brain section through the VTA, immunostained for TH. Black lines mark putative anatomical boundaries of the VTA. (B) Higher magnification view of the region outlined with a square in panel A, revealing TH-immunoreactivity filling the cytoplasm of the VTA cells (white arrows) and the segments of dendrites emanating from labeled cells. Scale bar = 400 µm in A, and 20 µm in B. (C) Volumetric density of TH-ir cells in WKY (black bars) and SHR (white bars) animals (n=3 WKY and 5 SHR; 4 sections per animal; p > 0.05 in Nested t-test). IP: interpeduncular nucleus; ml: medial lemniscus; SN: substantia nigra; Aq: aquaeductus cerebri.

**Figure 4.**
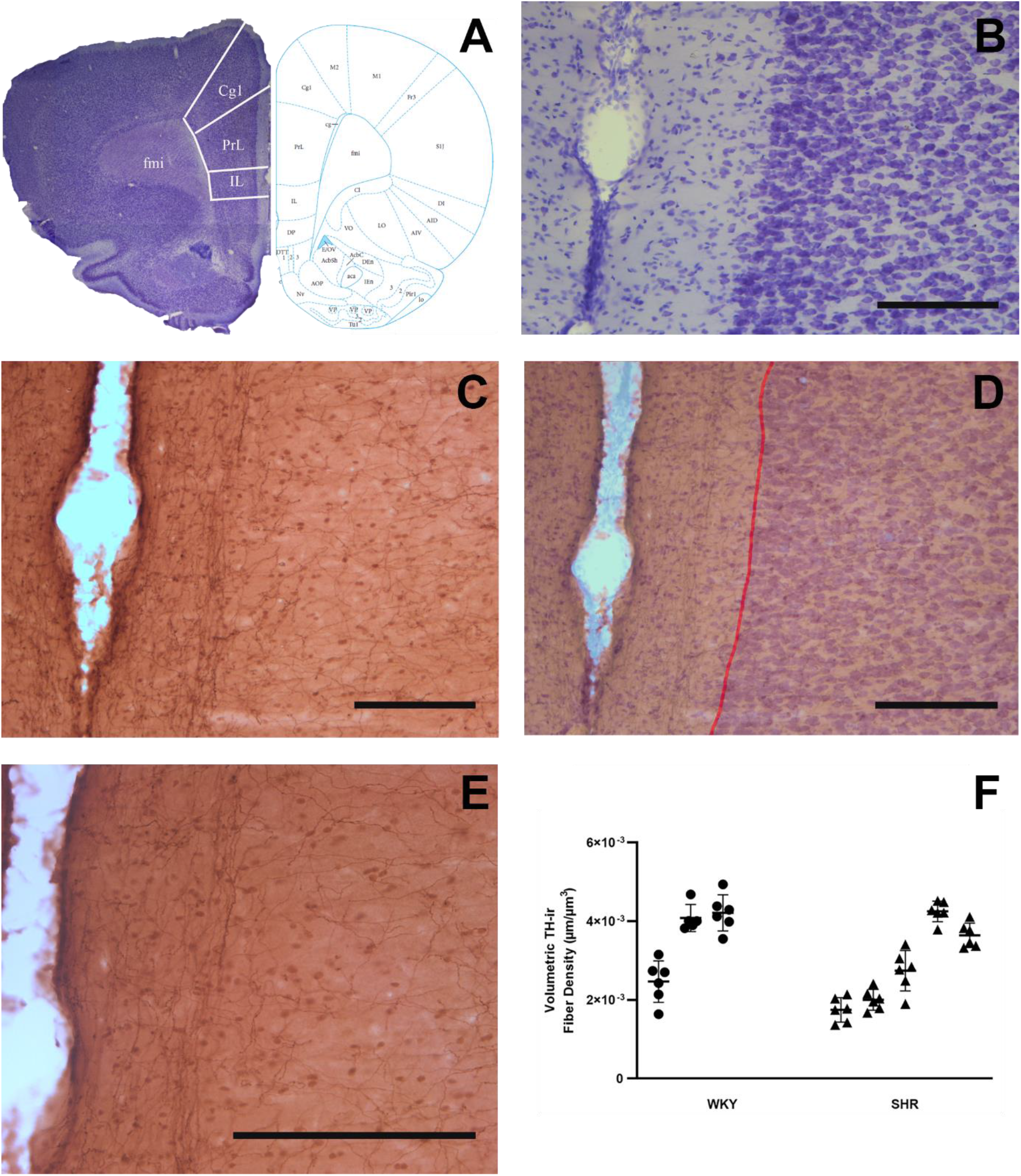
TH-ir fiber density in layer I of the PrL subregion of mPFC . (A-B) Coronal brain sections are used to identify the PrL of the mPFC (at around Bregma 3.00) on adjacent sections that are immunostained for TH (C and E). TH-ir fibers are encountered at all layers of the cortex, while those in Layer I are selectively oriented parallel to the pila surface (D-E). TH-ir fiber density measurements were confined in layer I, between the pial surface and Layers I and II border (red line). All scale bars = 250µm. (F) The volumetric density of fibers in Layer I of the mPFC-PrL of SHR and WKY rats (n=3 WKY and 5 SHR; 6 sections per animal; p>0.05 in Nested t-test). PrL: prelimbic area; IL: infralimbic area; Cg1: anterior cingulate cortex.

**Fig 5.**
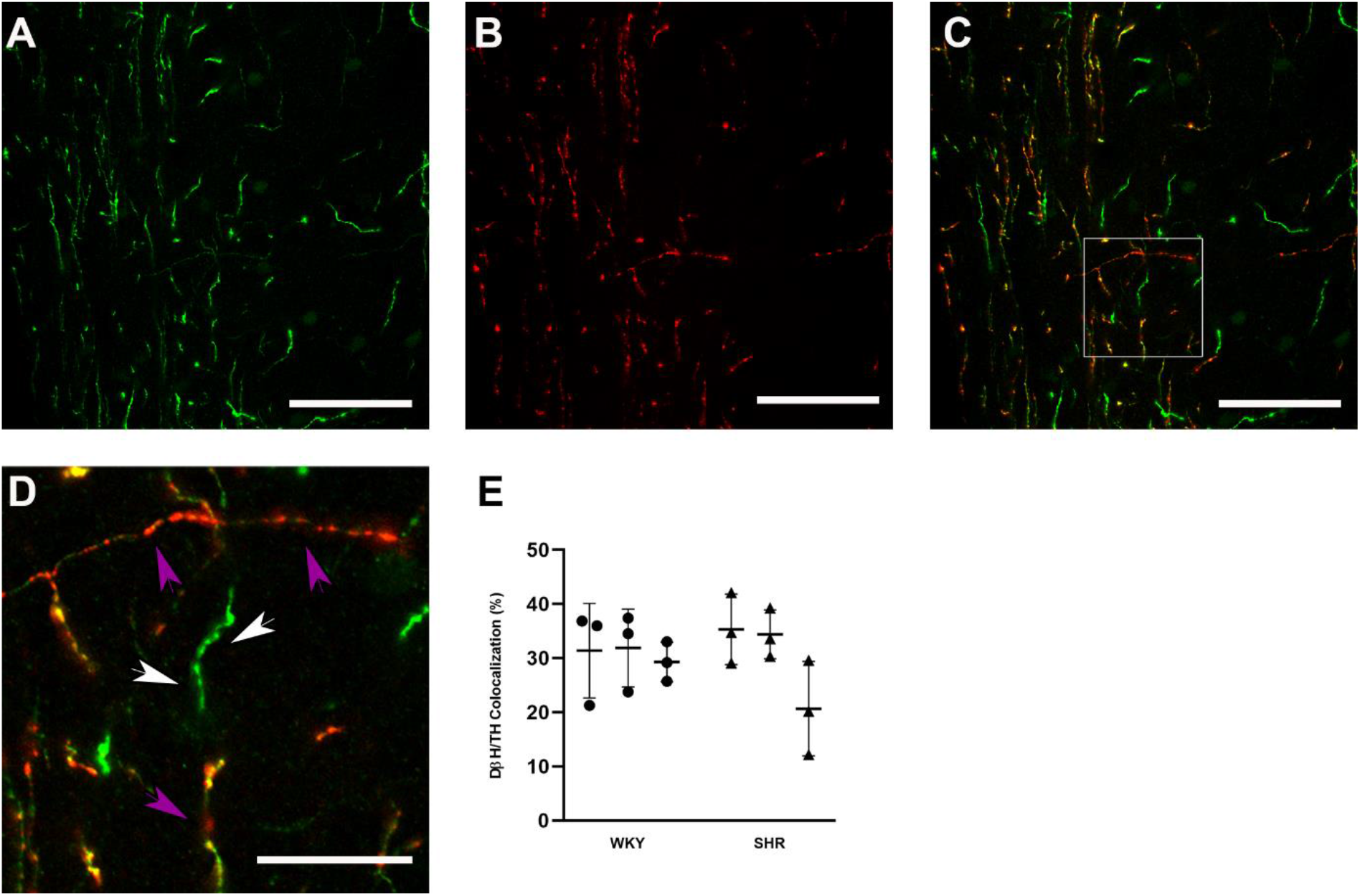
Colocalization analysis in layer I of the PrL subregion of mPFC. The axons immunolabeled for TH (green in A) and DBH (red in B) are imaged in the mPFC of a control brain. Merging of two images (C and D) reveals the presence of DBH (red; purple arrows) in TH (green; white arrows) axons and these dual labeled axons display patches of yellow fluorescence. Panel D is the high magnification of the area marked with a square frame in panel C. Note that all DBH-ir axons (purple arrows) are also TH-ir, but not all TH-ir (white arrows) are also DBH-ir. Scale bars: 50 µm in A-C and 15 µm in D. (E) The percentage of TH-ir axons in the mPFC. The measurements were obtained from 3 animals in each group 3 sections per animal; p>0.05 in the Nested t-test). No significant difference was found between the groups.

## 4. DISCUSSION

The result of the current study reveals that in SHR brains i) there is no dopaminergic cell loss in VTA, and ii) there is no reduction of DA axons in the PFC. These results invalidate our hypothesis and provide evidence that dopaminergic innervation in PFC cannot account for ADHD-like behavior observed in these animals.

Despite the high prevalence of ADHD, the etiology of this neurodevelopmental disorder has not been fully elucidated, yet (Thapar *et al*., 2009). Evidence supporting the role of DA in the pathophysiology of ADHD comes from studies in wide-ranging areas (Biederman & Faraone, 2002; Castellanos & Tannock, 2002; Swanson et al., 2007; Bowton et al., 2010). Nonetheless, studies on animal models aim at verifying the current postulate of a dysfunctional dopaminergic system in ADHD are scarce. The present results revealed the following: The mesocortical dopaminergic system appears to be normal in juvenile SHR, as suggested by (i) no alteration in the area density of TH-ir dopaminergic neurons in the VTA, (ii) no alterations in the volume density of TH-ir fibers in layer I of the PrL subregion of mPFC, (iii) no alteration in the percentage of TH-ir dopaminergic fibers in layer I of the PrL subregion of mPFC as revealed by subtraction of DBH-ir fibers from those that are TH-ir. To the best of our knowledge, this is the first study to evaluate the TH-ir neurons in VTA in the juvenile SHR to date. Similarly, the density of TH-positive fibers in the PRL in juvenile SHR and WKY was similar (Kozłowska *et al*., 2019).

Although SHR is the most widely used model for ADHD, hypertension can be confounding (Sagvolden, 2005). To eliminate this confounding factor, we measured SBP in SHR and WKY. Juvenile SHR exhibited no hypertension, furthermore, there were no differences between the strains with respect to SBP, as previously described (De Jong *et al*., 1995). We also tested the face validity of the SHR and their usual control, WKY, in an infrared beam-based activity meter to examine global motor activity to validate that symptoms of ADHD appear before SHR develops hypertension. In behavioral comparison in the activity meter, SHR displays an increased total distance traveled when compared to WKY. Previous work showed that SHR traveled greater total distances in open field tests than did WKY rats (Sagvolden et al., 2005, 2008, 2009; Tsai et al., 2017) which is interpreted as SHR rats present hyperactivity in motor function. These procedures have provided evidence that SHR mimics the basic behavioral characteristics of ADHD without having hypertension.

Since, TH is the rate-limiting enzyme of catecholamine biosynthesis, including DA, decreasing TH-ir density in mPFC may parallel reduced DA activity in the PFC, which produces hyperactivity in animals (Simon, 1981). Contrary to expectation, we observed no significant changes in the density of TH-ir dopaminergic fibers in mPFC in SHR compared to WKY. This result is in line with one of the earlier finding that TH-ir fiber density did not differ in SHR in the PrL and Cg1 subregions of the mPFC in a 5-week-old SHR (Kozłowska et al., 2019). Our results are also in accordance with a study by Leo et al., which observed significantly lower TH mRNA levels in the mesencephalon in SHR only at P5 and at P7 (Leo *et al*., 2003), which can explain the lack of differences in our study at the postnatal fourth week. The lack of differences in TH-ir fiber density between groups in this study should not rule out the involvement of the mesocortical system in ADHD, since we have not assessed the DA neurotransmission. Future studies will be needed to assess the expression of DAT and DA receptors, DA vesicular storage, DA transporter density, and DA availability at postsynaptic receptors in SHR to better understand the molecular mechanisms that cause dopaminergic hypofunction in ADHD.

ADHD is subclassified according to symptom clusters, hyperactive/impulsive, inattentive, or combined (American Psychiatric Association, 2022) which may have a heterogeneous origin. Interestingly, the hyperactive/impulsive subtype of ADHD may be qualitatively different from the ADHD inattentive subtype (Lahey al., 1994; Johansen et al., 2002). It has been suggested that the inattentive subtype may result from dysfunction in the inhibitory action of the frontal cortex, whereas the hyperactive/impulsive subtype may arise from an impairment of subcortical structures such as substantia nigra and striatum (Johansen et al., 2002; Solanto, 2002; Krause et al., 2003; Bowton et al., 2010; Miller et al., 2012). Substantia nigra plays an important role in movement and motor planning. The striatum serves as a mediator for many functions of the substantia nigra. The nigrostriatal dopaminergic pathway which projects from the substantia nigra to the striatum is closely linked with the striatum’s function (Nicola et al., 2000; Grillner & Mercuri, 2002). Hyperactivity in ADHD may result from excess dopaminergic activity in the striatum which is in line with increased striatal activity on PET in adolescents with ADHD relative to normal controls (Ernst *et al*., 1999). In our study, we found SHR displayed hyperactivity in the open field test, a result that is also similar to that of previous studies (Sagvolden et al., 2005, 2009; van den Bergh et al., 2006; Bayless et al., 2015; Tsai et al., 2017). The cause of locomotor hyperactivity in the SHR might be related to altered dopaminergic functioning in the striatum, but this will remain speculation without further research.

In the present experiment, SHR was more active than WKY in the open-field test. Although hyperactivity is necessary; it is not sufficient for an animal model of a combined subtype of ADHD. Previous studies reported that SHR did not show any impairment of attention-related behavior in the 5-choice serial reaction time task (van den Bergh *et al*., 2006) and a visual discrimination task (Thanos *et al*., 2010). We also showed no alteration in the TH-ir neurons in VTA and TH-ir fibers in mPFC in SHR. How should we interpret no alteration in the TH-ir neurons in VTA and TH-ir fibers in mPFC and the increased locomotor activity in the SHR? Perhaps, no alteration of the TH-ir fibers in the mPFC might be in line with previous findings that SHR does not display inattention symptoms of ADHD, and hyperactivity in the SHR might be a result of the dysregulation of dopamine in the striatum and SN. Even though the SHR is the most validated animal model of ADHD and shows some face validity, our data raises questions about the usefulness of the SHR as a model of the inattentive or combined subtype of ADHD. These results highlight that more research is required to further validate the use of SHRs as a suitable animal model for the inattentive or combined subtypes of ADHD as defined in DSM 5-TR. In this regard, previous findings should be treated with a degree of circumspection.

This study has limitations. First, as detailed characterization of the mesocortical dopaminergic fibers is not possible without classic anterograde or viral tracing, combining these techniques with multiple fluorescent immunohistochemistry would provide more precise anatomical information. Secondly, the use of different animals in immunohistochemistry and behavioral studies as those performed in different laboratories may prevent direct comparisons of the outcomes obtained from the two types of assessments. However, we used the same rat strains (SHR and WKY) from the same supplier (Charles River Laboratories) to obtain results that are directly comparable with each other. Thirdly, as substantia nigra and striatum are crucial parts of dopamine signaling and play a major role in motor control and attention, examining TH-ir neurons in the substantia nigra and TH-ir fibers in the striatum would provide in-depth information about the usefulness of the SHR as a suitable animal model for hyperactive/impulsive subtype of ADHD as defined in DSM-5-TR.

Overall, with these limitations in mind, the present study demonstrated that SHR exhibits more hyperactivity than WKY rats. Also, SHR was normotensive at postnatal 4-week age. There were no anatomical changes in the mesocortical dopaminergic system of SHR in comparison with age-matched WKY suggesting complex interaction of dopaminergic neurotransmission is not limited to anatomical changes. Future research is needed to disentangle the role of the mesocortical dopaminergic system in ADHD through other approaches.

## ACKNOWLEDGMENTS

We thank Alev Erisir for helpful discussions and comments, and training A.G.A. for procedures and data collection, Anila Tynan for her help with the immunohistochemical staining, Amy Adams for her help in animal care, and Mert Can Aydin (B.A, Middle East Technical University) for help with the figures.

## AUTHOR CONTRIBUTIONS

A.G.A. Designed the study, performed the experiments, collected, and analyzed the data, wrote the original draft, reviewed & edited the manuscript, visualization, project administration, and funding acquisition. E.A. Designed the study, analyzed the data, reviewed & edited the manuscript, visualization, project administration, funding acquisition, and supervision. Both authors approved the final version of the manuscript.

## DATA ACCESSIBILITY

Data can be obtained by contacting the corresponding author.

## ABBREVIATIONS

ADHD: attention deficit hyperactivity disorder
VTA: ventral tegmental area
DA: dopamine
NE: norepinephrine
PFC: prefrontal cortex
mPFC: medial prefrontal cortex
PrL: prelimbic area
SHR: spontaneously hypertensive rat
WKY: wistar kyoto rat
TH: tyrosine hydroxylase
DBH: dopamine β-hydroxylase

## Graphical Abstract

Immunohistochemistry analysis showed that there is an alteration in TH-ir dopaminergic neurons in VTA and TH-ir fibers in PFC in juvenile SHR. Although the mesocortical dopaminergic system appears to be normal, SHR showed increased locomotor activity than WKY in the open field test.

## REFERENCES

American Psychiatric Association (2013) Diagnostic and Statistical Manual of Mental Disorders. In: (DSM-5). American Psychiatric Publishing.

American Psychiatric Association (2022) Diagnostic and Statistical Manual of Mental Disorders (DSM-5-TR). American Psychiatric Association Publishing.

Arnsten, A.F.T. (2011) Catecholamine influences on dorsolateral prefrontal cortical networks. Biol Psychiatry, 69, 89–99.

Barkley, R.A. (1990) A critique of current diagnostic criteria for attention deficit hyperactivity disorder: Clinical and research implications. Journal of Developmental and Behavioral Pediatrics, 11, 343–352.

Bayer, V.E. & Pickel, V.M. (1990) Ultrastructural localization of tyrosine hydroxylase in the rat ventral tegmental area: Relationship between immunolabeling density and neuronal associations. Journal of Neuroscience, 10, 2996–3013.

Bayless, D.W., Perez, M.C., & Daniel, J.M. (2015) Comparison of the validity of the use of the spontaneously hypertensive rat as a model of attention deficit hyperactivity disorder in males and females. Behavioural Brain Research, 286, 85–92.

Berger, D.F. & Sagvolden, T. (1998) Sex differences in operant discrimination behaviour in an animal model of attention-deficit hyperactivity disorder. Behavioural Brain Research, 94, 73–82.

Biederman, J. (2005) Attention-deficit/hyperactivity disorder: a selective overview. Biol Psychiatry, 57, 1215–1220.

Biederman, J. & Faraone, S. V. (2002) Current concepts on the neurobiology of Attention-Deficit/Hyperactivity Disorder. J Atten Disord,.

Biederman, J., Milberger, S., Faraone, S. V., Kiely, K., Guite, J., Mick, E., Ablon, S., Warburton, R., & Reed, E. (1995) Family-Environment Risk Factors for Attention-Deficit Hyperactivity Disorder: A Test of Rutter’s Indicators of Adversity. Arch Gen Psychiatry, 52, 464–470.

Botanas, C.J., Lee, H., de la Peña, J.B., dela Peña, I.J., Woo, T., Kim, H.J., Han, D.H., Kim, B.N., & Cheong, J.H. (2016) Rearing in an enriched environment attenuated hyperactivity and inattention in the Spontaneously Hypertensive Rats, an animal model of Attention-Deficit Hyperactivity Disorder. Physiol Behav, 155, 30–37.

Bowton, E., Saunders, C., Erreger, K., Sakrikar, D., Matthies, H.J., Sen, N., Jessen, T., Colbran, R.J., Caron, M.G., Javitch, J.A., Blakely, R.D., & Galli, A. (2010) Dysregulation of dopamine transporters via dopamine D2 autoreceptors triggers anomalous dopamine efflux associated with attention-deficit hyperactivity disorder. Journal of Neuroscience, 30, 6048–6057.

Brozoski, T.J., Brown, R.M., Rosvold, H.E., & Goldman, P.S. (1979) Cognitive deficit caused by regional depletion of dopamine in prefrontal cortex of rhesus monkey. Science (1979), 205, 929–932.

Bubser, M. & Schmidt, W.J. (1990) 6-Hydroxydopamine lesion of the rat prefrontal cortex increases locomotor activity, impairs acquisition of delayed alternation tasks, but does not affect uninterrupted tasks in the radial maze. Behavioural Brain Research, 37, 157–168.

Canales, J.J. & Iversen, S.D. (1998) Behavioural topography in the striatum: Differential effects of quinpirole and D-amphetamine microinjections. Eur J Pharmacol, 362, 111–119.

Carr, G.D. & White, N.M. (1987) Effects of systemic and intracranial amphetamine injections on behavior in the open field: A detailed analysis. Pharmacol Biochem Behav, 27, 113–122.

Castellanos, F.X. & Tannock, R. (2002) Neuroscience of attention-deficit/hyperactivity disorder: The search for endophenotypes. Nat Rev Neurosci,.

Castner, S.A. & Williams, G. V. (2007) Tuning the engine of cognition: A focus on NMDA/D1 receptor interactions in prefrontal cortex. Brain Cogn, 63, 94–122.

Cheng, J., Liu, A., Shi, M.Y., & Yan, Z. (2017) Disrupted glutamatergic transmission in prefrontal cortex contributes to behavioral abnormality in an animal model of ADHD. Neuropsychopharmacology, 42, 2096–2104.

Cook, E.H., Stein, M.A., Krasowski, M.D., Cox, N.J., Olkon, D.M., Kieffer, J.E., & Leventhal, B.L. (1995) Association of attention-deficit disorder and the dopamine transporter gene. Am J Hum Genet, 56, 993–998.

Cortese, S., Kelly, C., Chabernaud, C., Proal, E., Di Martino, A., Milham, M.P., & Castellanos, F.X. (2012) Toward systems neuroscience of ADHD: A meta-analysis of 55 fMRI sudies. American Journal of Psychiatry, 169, 1038–1055.

De Jong, W., Linthorst, A.C.E., & Versteeg, H.G. (1995) The nigrostriatal dopamine system and the development of hypertension in the spontaneously hypertensive rat. Arch Mal Coeur Vaiss, 88, 1193–1196.

Ernst, M., Zametkin, A.J., Matochik, J.A., Pascualvaca, D., Jons, P.H., & Cohen, R.M. (1999) High Midbrain [_18_F] DOPA Accumulation in Children With Attention Deficit Hyperactivity Disorder. American Journal of Psychiatry, 156, 1209–1215.

Evenden, J.L. (1998) Serotonergic and steroidal influences on impulsive behaviour in rats.

Faraone, S. V, Sergeant, J., Gillberg, C., & Biederman, J. (2003) The worldwide prevalence of ADHD: is it an American condition? World Psychiatry, 2, 104–113.

Fernández, A., Quintero, J., Hornero, R., Zuluaga, P., Navas, M., Gómez, C., Escudero, J., García-Campos, N., Biederman, J., & Ortiz, T. (2009) Complexity Analysis of Spontaneous Brain Activity in Attention-Deficit/Hyperactivity Disorder: Diagnostic Implications. Biol Psychiatry, 65, 571–577.

Floresco, S.B. & Magyar, O. (2006) Mesocortical dopamine modulation of executive functions: Beyond working memory. Psychopharmacology (Berl), 188, 567–585.

Giedd, J.N., Blumenthal, J., Molloy, E., & Castellanos, F.X. (2001) Brain imaging of attention deficit/hyperactivity disorder. Ann N Y Acad Sci, 931, 33–49.

Granon, S., Passetti, F., Thomas, K.L., Dalley, J.W., Everitt, B.J., & Robbins, T.W. (2000) Enhanced and impaired attentional performance after infusion of D1 dopaminergic receptor agents into rat prefrontal cortex. Journal of Neuroscience, 20, 1208–1215.

Grillner, P. & Mercuri, N.B. (2002) Intrinsic membrane properties and synaptic inputs regulating the firing activity of the dopamine neurons. Behavioural Brain Research, 130, 149–169.

Gundersen, H.J.G. (1977) Notes on the estimation of the numerical density of arbitrary profiles: the edge effect. J Microsc, 111, 219–223.

Gundersen, H.J.G. & Jensen, E.B. (1987) The efficiency of systematic sampling in stereology and its prediction*. J Microsc, 147, 229–263.

Harvey, R.C., Jordan, C.J., Tassin, D.H., Moody, K.R., Dwoskin, L.P., & Kantak, K.M. (2013) Performance on a strategy set shifting task during adolescence in a genetic model of attention deficit/hyperactivity disorder: Methylphenidate vs. atomoxetine treatments. Behavioural Brain Research, 244, 38–47.

Heidbreder, C.A. & Groenewegen, H.J. (2003) The medial prefrontal cortex in the rat: Evidence for a dorso-ventral distinction based upon functional and anatomical characteristics. Neurosci Biobehav Rev, 27, 555–579.

Himelstein, J., Newcorn, J.H., & Halperin, J.M. (2000) The neurobiology of attention-deficit hyperactivity disorder. Front Biosci, 5.

Hokfelt, T., Johansson, O., Fuxe, K., Goldstein, M., & Park, D. (1976) Immunohistochemical studies on the localization and distribution of monoamine neuron systems in the rat brain. I. Tyrosine hydroxylase in the mes and diencephalon. Med Biol, 54, 427–453.

Johansen, E.B., Aase, H., Meyer, A., & Sagvolden, T. (2002) Attention-deficit/hyperactivity disorder (ADHD) behaviour explained by dysfunctioning reinforcement and extinction processes. Behavioural Brain Research, 130, 37–45.

Jucaite, A., Fernell, E., Halldin, C., Forssberg, H., & Farde, L. (2005) Reduced midbrain dopamine transporter binding in male adolescents with attention-deficit/hyperactivity disorder: Association between striatal dopamine markers and motor hyperactivity. Biol Psychiatry, 57, 229–238.

Kantak, K.M., Singh, T., Kerstetter, K.A., Dembro, K.A., Mutebi, M.M., Harvey, R.C., Deschepper, C.F., & Dwoskin, L.P. (2008) Advancing the spontaneous hypertensive rat model of attention deficit/hyperactivity disorder. Behavioral Neuroscience, 122, 340–357.

Kelly, A.M.C., Margulies, D.S., & Castellanos, F.X. (2007) Recent advances in structural and functional brain imaging studies of attention-deficit/hyperactivity disorder. Curr Psychiatry Rep, 9, 401–407.

Kessler, R.C., Adler, L., Berkley, R., Biederman, J., Conners, C.K., Demler, O., Faraone, S. V., Greenhill, L.L., Howes, M.J., Secnik, K., Spencer, T., Ustun, T.B., Walters, E.E., & Zaslavsky, A.M. (2006) The prevalence and correlates of adult ADHD in the United States: Results from the National Comorbidity Survey Replication. American Journal of Psychiatry, 163, 716–723.

Kozłowska, A., Wojtacha, P., Równiak, M., Kolenkiewicz, M., & Huang, A.C.W. (2019) ADHD pathogenesis in the immune, endocrine and nervous systems of juvenile and maturating SHR and WKY rats. Psychopharmacology (Berl), 236, 2937–2958.

Krause, K.-H., Dresel, S.H., Krause, J., la Fougere, C., & Ackenheil, M. (2003) The dopamine transporter and neuroimaging in attention deficit hyperactivity disorder. Neurosci Biobehav Rev, 27, 605–613.

Kreppner, J.M., O’Connor, T.G., Rutter, M., Beckett, C., Castle, J., Croft, C., Dunn, J., & Groothues, C. (2001) Can inattention/overactivity be an institutional deprivation syndrome? J Abnorm Child Psychol, 29, 513–528.

Lahey B B, A.B.M.K.B.J.G.L.H.G. & W., B.R.A., N.J.J.B.K.L.F.P.J., O.T.P.D.H.E.L. (1994) DSM-IV field trials for attention deficit hyperactivity disorder in children and adolescents. American Journal of Psychiatry, 151, 1673–1685.

Lehnen, A.M., Leguisamo, N.M., Pinto, G.H., Markoski, M.M., de Angelis, K., Machado, U.F., & Schaan, B. (2010) The beneficial effects of exercise in rodents are preserved after detraining: a phenomenon unrelated to GLUT4 expression. Cardiovasc Diabetol, 9, 67.

Leo, D., Sorrentino, E., Volpicelli, F., Eyman, M., Greco, D., Viggiano, D., Di Porzio, U., & Perrone-Capano, C. (2003) Altered midbrain dopaminergic neurotransmission during development in an animal model of ADHD. Neurosci Biobehav Rev, 27, 661–669.

Ludolph, A.G., Kassubek, J., Schmeck, K., Glaser, C., Wunderlich, A., Buck, A.K., Reske, S.N., Fegert, J.M., & Mottaghy, F.M. (2008) Dopaminergic dysfunction in attention deficit hyperactivity disorder (ADHD), differences between pharmacologically treated and never treated young adults: A 3,4-dihdroxy-6-[18F]fluorophenyl-l-alanine PET study. Neuroimage, 41, 718–727.

Mick, E., Biederman, J., Faraone, S. V., Sayer, J., & Kleinman, S. (2002) Case-Control Study of Attention-Deficit Hyperactivity Disorder and Maternal Smoking, Alcohol Use, and Drug Use during Pregnancy. J Am Acad Child Adolesc Psychiatry, 41, 378–385.

Mill, J., Sagvolden, T., & Asherson, P. (2005) Sequence analysis of Drd2, Drd4, and Dat1 in SHR and WKY rat strains. Behavioral and Brain Functions, 1, 24.

Miller, E.M., Pomerleau, F., Huettl, P., Russell, V.A., Gerhardt, G.A., & Glaser, P.E.A. (2012) The spontaneously hypertensive and Wistar Kyoto rat models of ADHD exhibit sub-regional differences in dopamine release and uptake in the striatum and nucleus accumbens. Neuropharmacology, 63, 1327–1334.

Mook, D.M., Jeffrey, J., & Neuringer, A. (1993) Spontaneously hypertensive rats (SHR) readily learn to vary but not repeat instrumental responses. Behav Neural Biol, 59, 126–135.

Moon, S.J., Kim, C.J., Lee, Y.J., Hong, M., Han, J., & Bahn, G.H. (2014) Effect of atomoxetine on hyperactivity in an animal model of attention-deficit/hyperactivity disorder (ADHD). PLoS One, 9.

Morrison, J.H., Grzanna, R., Molliver, M.E., & Coyle, J.T. (1978) The distribution and orientation of noradrenergic fibers in neocortex of the rat: An immunofluorescence study. Journal of Comparative Neurology, 181, 17–39.

Murphy, B.L., Arnsten, A.F.T., Goldman-Rakic, P.S., & Roth, R.H. (1996) Increased dopamine turnover in the prefrontal cortex impairs spatial working memory performance in rats and monkeys. Proc Natl Acad Sci U S A, 93, 1325–1329.

Naneix, F., Marchand, A.R., Scala, G.D., Pape, J.-R., & Coutureau, E. (2009) A Role for Medial Prefrontal Dopaminergic Innervation in Instrumental Conditioning. Journal of Neuroscience, 29, 6599–6606.

Nicola, S.M., Surmeier, D.J., & Malenka, R.C. (2000) Dopaminergic Modulation of Neuronal Excitability in the Striatum and Nucleus Accumbens. Annu Rev Neurosci, 23, 185–215.

Nigg, J.T. & Casey, B.J. (2005) An integrative theory of attention-deficit/hyperactivity disorder based on the cognitive and affective neurosciences. Dev Psychopathol, 17, 785–806.

Papa, M., Diewald, L., Carey, M.P., Esposito, F.J., Gironi Carnevale, U.A., & Sadile, A.G. (2002) A rostro-caudal dissociation in the dorsal and ventral striatum of the juvenile SHR suggests an anterior hypo- and a posterior hyperfunctioning mesocorticolimbic system. Behavioural Brain Research, 130, 171–179.

Paxinos, G. & Watson, C. (2007) The Rat Brain in Stereotaxic Coordinates 6th Edition, 6th edn. Academic Press.

Pickel, V.M., Joh, T.H., & Reis, D.J. (1976) Monoamine synthesizing enzymes in central dopaminergic, noradrenergic and serotonergic neurons. Immunocytochemical localization by light and electron microscopy. Journal of Histochemistry and Cytochemistry, 24, 792–806.

Pirot, S., Godbout, R., Mantz, J., Tassin, J.P., Glowinski, J., & Thierry, A.M. (1992) Inhibitory effects of ventral tegmental area stimulation on the activity of prefrontal cortical neurons: Evidence for the involvement of both dopaminergic and GABAergic components. Neuroscience, 49, 857–865.

Polanczyk, G., De Lima, M.S., Horta, B.L., Biederman, J., & Rohde, L.A. (2007) The worldwide prevalence of ADHD: A systematic review and metaregression analysis. American Journal of Psychiatry, 164, 942–948.

Prince, J. (2008) Catecholamine dysfunction in attention-deficit/hyperactivity disorder an update. J Clin Psychopharmacol, 28, 39–45.

Qin, L., Liu, W., Ma, K., Wei, J., Zhong, P., Cho, K., & Yan, Z. (2016) The ADHD-linked human dopamine D4 receptor variant D4.7 induces over-suppression of NMDA receptor function in prefrontal cortex. Neurobiol Dis, 95, 194–203.

Robbins, T.W. & Arnsten, A.F.T. (2009) The Neuropsychopharmacology of Fronto-Executive Function: Monoaminergic Modulation. Annu Rev Neurosci, 32, 267–287.

Russell, V., de Villiers, A., Sagvolden, T., Lamm, M., & Taljaard, J. (1998) Differences between electrically-, ritalin- and D-amphetamine-stimulated release of [3H]dopamine from brain slices suggest impaired vesicular storage of dopamine in an animal model of Attention-Deficit Hyperactivity Disorder. Behavioural Brain Research, 94, 163–171.

Sagvolden, T. (2000) Behavioral validation of the spontaneously hypertensive rat (SHR) as an animal model of attention-deficit/hyperactivity disorder (AD/HD). Neurosci Biobehav Rev, 24, 31–39.

Sagvolden, T., Russell, V.A., Aase, H., Johansen, E.B., & Farshbaf, M. (2005) Rodent models of attention-deficit/hyperactivity disorder. Biol Psychiatry, 57, 1239–1247.

Sagvolden, T., DasBanerjee, T., Zhang-James, Y., Middleton, F.A., & Faraone, S. V. (2008) Behavioral and genetic evidence for a novel animal model of Attention-Deficit/Hyperactivity Disorder Predominantly Inattentive Subtype. Behav Brain Funct, 4, 56.

Sagvolden, T. & Johansen, E.B. (2012) Rat models of ADHD. Curr Top Behav Neurosci, 9, 301–315.

Sagvolden, T., Johansen, E.B., Wøien, G., Walaas, S.I., Storm-Mathisen, J., Bergersen, L.H., Hvalby, Ø., Jensen, V., Aase, H., Russell, V.A., Killeen, P.R., DasBanerjee, T., Middleton, F.A., & Faraone, S. V. (2009) The spontaneously hypertensive rat model of ADHD - The importance of selecting the appropriate reference strain. Neuropharmacology, 57, 619–626.

Sawaguchi, T. & Goldman-Rakic, P.S. (1994) The role of D1-dopamine receptor in working memory: Local injections of dopamine antagonists into the prefrontal cortex of rhesus monkeys performing an oculomotor delayed-response task. J Neurophysiol, 71, 515–528.

Sesack, S.R. & Bunney, B.S. (1989) Pharmacological characterization of the receptor mediating electrophysiological responses to dopamine in the rat medial prefrontal cortex: A microiontophoretic study. Journal of Pharmacology and Experimental Therapeutics, 248, 1323–1333.

Sesack, S.R., Hawrylak, V.A., Matus, C., Guido, M.A., & Levey, A.I. (1998) Dopamine axon varicosities in the prelimbic division of the rat prefrontal cortex exhibit sparse immunoreactivity for the dopamine transporter. Journal of Neuroscience, 18, 2697–2708.

Simon, H. (1981) Dopaminergic A10 neurons and frontal system. J Physiol (Paris), 77, 81–95.

Smith, A.B., Taylor, E., Brammer, M., Halari, R., & Rubia, K. (2008) Reduced activation in right lateral prefrontal cortex and anterior cingulate gyrus in medication-naïve adolescents with attention deficit hyperactivity disorder during time discrimination. J Child Psychol Psychiatry, 49, 977–985.

Solanto, M. V (2002) Dopamine dysfunction in AD/HD: integrating clinical and basic neuroscience research. Behavioural Brain Research, 130, 65–71.

Somkuwar, S.S., Kantak, K.M., Bardo, M.T., & Dwoskin, L.P. (2016) Adolescent methylphenidate treatment differentially alters adult impulsivity and hyperactivity in the Spontaneously Hypertensive Rat model of ADHD. Pharmacol Biochem Behav, 141, 66–77.

Sterio, D.C. (1984) The unbiased estimation of number and sizes of arbitrary particles using the disector. J Microsc, 134, 127–136.

Sullivan, R.M. & Brake, W.G. (2003) What the rodent prefrontal cortex can teach us about attention-deficit/ hyperactivity disorder: The critical role of early developmental events on prefrontal function. Behavioural Brain Research, 146, 43–55.

Swanson, J.M., Kinsbourne, M., Nigg, J., Lanphear, B., Stefanatos, G.A., Volkow, N., Taylor, E., Casey, B.J., Castellanos, F.X., & Wadhwa, P.D. (2007) Etiologic subtypes of attention-deficit/hyperactivity disorder: Brain imaging, molecular genetic and environmental factors and the dopamine hypothesis. Neuropsychol Rev, 17, 39–59.

Swanson, J.M., Sergeant, J.A., Taylor, E., Sonuga-Barke, E.J.S., Jensen, P.S., & Cantwell, D.P. (2001) Attention-deficit hyperactivity disorder and hyperkinetic disorder. Lancet, 351, 39–43.

Thanos, P.K., Ivanov, I., Robinson, J.K., Michaelides, M., Wang, G.-J., Swanson, J.M., Newcorn, J.H., & Volkow, N.D. (2010) Dissociation between spontaneously hypertensive (SHR) and Wistar–Kyoto (WKY) rats in baseline performance and methylphenidate response on measures of attention, impulsivity and hyperactivity in a Visual Stimulus Position Discrimination Task. Pharmacol Biochem Behav, 94, 374–379.

Thapar, A., Rice, F., Hay, D., Boivin, J., Langley, K., Van Den Bree, M., Rutter, M., & Harold, G. (2009) Prenatal smoking might not cause attention-deficit/hyperactivity disorder: Evidence from a novel design. Biol Psychiatry, 66, 722–727.

Tsai, M.L., Kozłowska, A., Li, Y.S., Shen, W.L., & Huang, A.C.W. (2017) Social factors affect motor and anxiety behaviors in the animal model of attention-deficit hyperactivity disorders: A housing-style factor. Psychiatry Res, 254, 290–300.

van den Bergh, F.S., Bloemarts, E., Chan, J.S.W., Groenink, L., Olivier, B., & Oosting, R.S. (2006) Spontaneously hypertensive rats do not predict symptoms of attention-deficit hyperactivity disorder. Pharmacol Biochem Behav, 83, 380–390.

Volkow, N.D., Wang, G.J., Kollins, S.H., Wigal, T.L., Newcorn, J.H., Telang, F., Fowler, J.S., Zhu, W., Logan, J., Ma, Y., Pradhan, K., Wong, C., & Swanson, J.M. (2009) Evaluating dopamine reward pathway in ADHD: Clinical Implications. JAMA - Journal of the American Medical Association, 302, 1084–1091.

Volkow, N.D., Wang, G.J., Newcorn, J., Telang, F., Solanto, M. V., Fowler, J.S., Logan, J., Ma, Y., Schulz, K., Pradhan, K., Wong, C., & Swanson, J.M. (2007) Depressed dopamine activity in caudate and preliminary evidence of limbic involvement in adults with attention-deficit/hyperactivity disorder. Arch Gen Psychiatry, 64, 932–940.

West, M.J., Slomianka, L., & Gundersen, H.J.G. (1991) Unbiased stereological estimation of the total number of neurons in the subdivisions of the rat hippocampus using the optical fractionator. Anat Rec, 231, 482–497.

Witelson, S.F., Glezer, I.I., & Kigar, D.L. (1995) Women have greater density of neurons in posterior temporal cortex. Journal of Neuroscience, 15, 3418–3428.

Xing, B., Li, Y.C., & Gao, W.J. (2016) Norepinephrine versus dopamine and their interaction in modulating synaptic function in the prefrontal cortex. Brain Res, 1641, 217–233.

Zahrt, J., Taylor, J.R., Mathew, R.G., & Arnsten, A.F.T. (1997) Supranormal stimulation of D1 dopamine receptors in the rodent prefrontal cortex impairs spatial working memory performance. Journal of Neuroscience, 17, 8528–8535.

